# Mechanofiltration Enables High-Throughput Measurements of Bacterial Cell Mechanics

**DOI:** 10.64898/2026.09.01.748625

**Authors:** Kelsey G. DeFrates, Jae Won Hwang, Gissell Jimenez, Kerwyn Casey Huang, Joanne N. Engel, Christopher J. Hernandez

**Affiliations:** Department of Orthopaedic Surgery, University of California, San Francisco, California, United States; Department of Bioengineering, Stanford University, Stanford, California, United States; Department of Microbiology & Immunology, Stanford University School of Medicine, Stanford, California, United States; Biohub, San Francisco, California, United States; Department of Medicine, University of California, San Francisco, San Francisco, California, United States; Department of Microbiology and Immunology, University of California, San Francisco, San Francisco, California, United States; Department of Bioengineering and Therapeutic Sciences, University of California, San Francisco, San Francisco, California, United States; Department of Bioengineering, University of California, Berkeley, Berkeley, California, United States

**Keywords:** bacteria, biomechanics, cell stiffness, high-throughput screening, centrifugation

## Abstract

Bacteria experience diverse mechanical forces throughout their natural environments, yet quantitative measurements of bacterial biomechanics remain challenging because most existing techniques require specialized instrumentation, such as atomic force microscopy (AFM) or microfluidic devices. Here, we introduce mechanofiltration, a simple, high-throughput assay that estimates whole-cell mechanics using standard laboratory equipment. In mechanofiltration, bacterial suspensions are centrifuged through porous membranes in multiwell plates. Pressure generated during centrifugation drives cells toward pores that are smaller than cell width, requiring cells to deform to transit through the membrane. By combining recovery on the opposite side of the filter with measurements of cell size, the assay estimates maximum cell deformation, a size-corrected proxy for whole-cell stiffness. We show that mechanofiltration detects established reductions in *Escherichia coli* cell stiffness caused by genetic disruption of load-bearing cell envelope components, treatment with outer membrane– destabilizing agents, and sublethal exposure to antibiotics. Fold changes in maximum deformation closely agree with published measurements obtained using AFM, cell-bending assays, and osmotic shock experiments. Mechanofiltration also reproduces previously reported mechanical differences among bacterial species with distinct morphologies and envelope architectures. Together, these results establish mechanofiltration as an accessible, inexpensive, and scalable approach for identifying genetic and environmental determinants of bacterial mechanics and for high-throughput screening of bacterial biomechanical phenotypes.

## INTRODUCTION

Biomechanical properties regulate diverse aspects of cellular physiology and function. In mammals, changes in cell mechanics accompany cell-cycle progression^1,2^, differentiation^3–5^, migration^6,7^, immune activation^8–10^, and disease progression^11–14^. Mechanical properties are equally important in bacteria, where they contribute to cell-shape maintenance during growth and division, regulate adaptation to osmotic stress^15–19^, and control migration through constricted environments, such as porous soils or microscale cavities within host tissues and engineered materials^20–25^. Mechanical deformation also serves as a biological signal by activating stress response pathways, regulating mechanosensitive ion channels, and influencing the assembly of trans-envelope protein complexes^26–30^. Together, these observations highlight bacterial mechanics as an important determinant of microbial physiology, adaptation, and survival.

Despite their biological importance, the mechanical properties of bacteria are not routinely measured because the small size of bacterial cells (∼1 µm) makes it challenging to apply controlled mechanical loads and quantify deformation at the single-cell level. Atomic force microscopy (AFM), which measures cantilever deflection during contact with the cell surface, is among the most widely used techniques for studying bacterial mechanics. AFM can quantify whole-cell stiffness or, when combined with mechanical modeling, infer material properties of individual cell-envelope components^31–35^. However, AFM is inherently low throughput, requires immobilization of cells, and is sensitive to environmental conditions^36,37^. Other approaches have examined bacterial mechanics by measuring deformation during osmotic shock, cell bending under fluid flow or optical trapping^38,39^, and cell transport through tapered microfluidic nanochannels under applied pressure^40,41^. Although these methods overcome some limitations of AFM, they generally require specialized instrumentation, high-resolution microscopy, or microfabricated devices, limiting their accessibility and throughput for large-scale studies.

Currently, only one approach enables high-throughput measurements of bacterial biomechanics. In this method, bacterial growth is monitored within agarose gels of defined stiffness^37,42,43^, where compressive forces generated by the surrounding matrix slow cell elongation in proportion to whole-cell stiffness^37,42,43^. This strategy has enabled large-scale biomechanical screens of model organisms^42,43^. However, because the applied mechanical load arises from cell growth itself, changes in growth physiology and mechanics are intrinsically coupled^44–48^. Applying externally controlled mechanical forces to bacteria could provide a more direct and broadly applicable approach for high-throughput measurements of bacterial mechanics.

Using microfluidic devices, we previously demonstrated that whole-cell stiffness governs the ability of bacteria to deform through nanoscale constrictions under fluid pressure^40,41^. Although this study provided quantitative measurements of bacterial mechanics, our microfluidic devices demand specialized fabrication and substantial technical expertise, and only characterize a relatively small number of strains or species per experiment, require, and. To overcome these limitations, here, we introduce mechanofiltration, a simple, multiwell assay that estimates whole-cell mechanics by quantifying bacterial transport through porous membranes during centrifugation. We show that mechanofiltration detects established genetic and chemical perturbations of bacterial mechanics, reproduces previously reported mechanical differences among bacterial species, and can be implemented using standard laboratory equipment without specialized instrumentation or complex mechanical modeling. Mechanofiltration therefore provides an accessible, high-throughput platform for identifying genetic and environmental determinants of bacterial biomechanics.

## RESULTS

### Development and Validation of the Mechanofiltration Assay

In mechanofiltration, hydrostatic pressure pushes bacteria toward a filter with pores smaller than the cell width (**Figure 1**). Because softer cells are expected to undergo greater deformation under a given loading condition, differences in cell stiffness can be inferred from the fraction of cells that pass through the filter after accounting for variation in cell size. To enable rapid analysis in a multiwell format, mechanofiltration can be performed using porous transwell inserts. In a typical experiment, a bacterial suspension, termed the stock, is added to the apical chamber above the filter, and the plate is centrifuged. Centrifugation generates a pressure difference across the filter that drives liquid from the apical chamber into the basal chamber. Again, cells with widths greater than the pore diameter must deform to also pass through the filter and enter what is referred to as the permeate. Viable cells in the stock and permeate are subsequently quantified using standard microbiological methods, such as colony counting or optical density.

**Figure 1.**
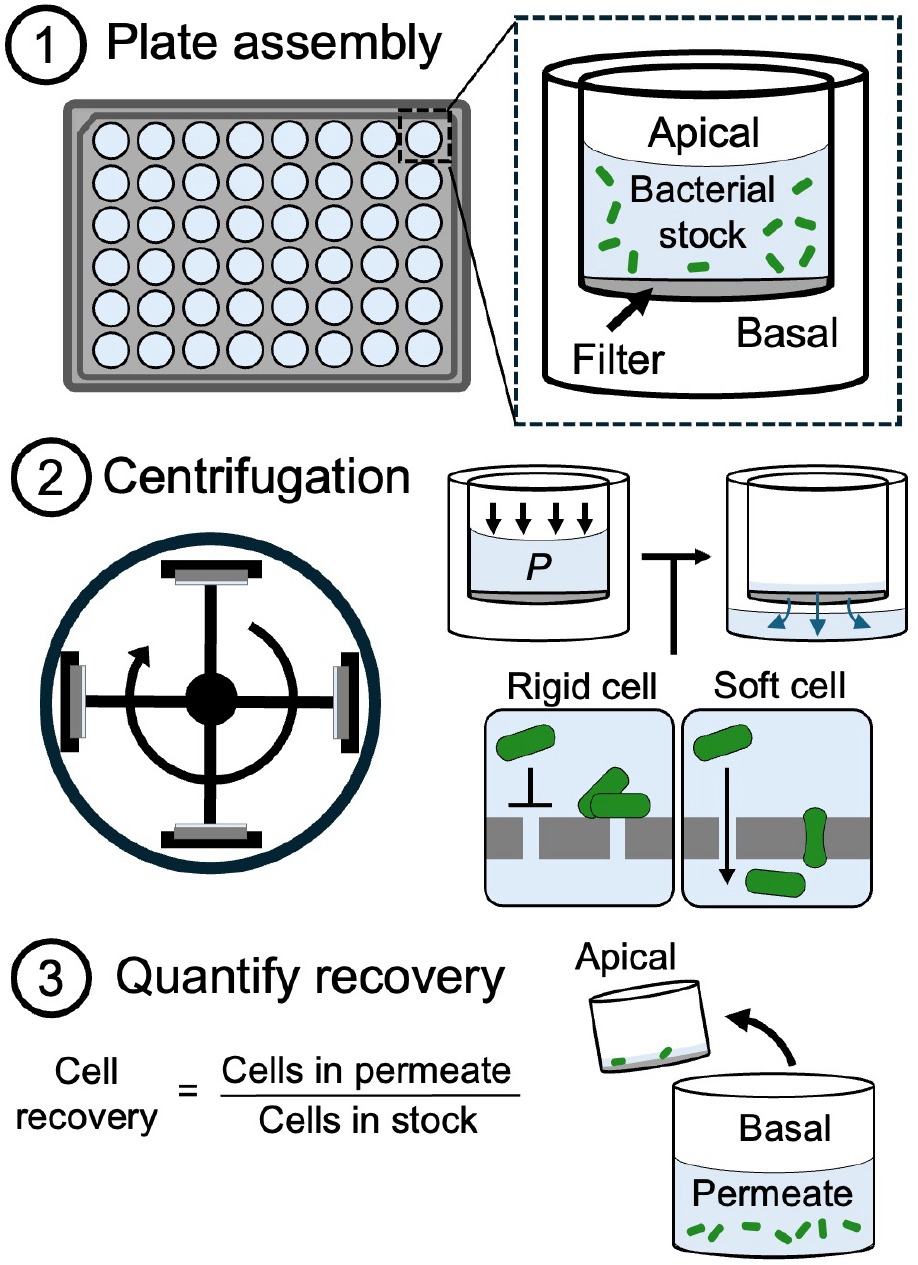
Overview of the mechanofiltration assay. (1) A bacterial suspension (stock) is added to the apical compartment of porous transwell inserts in a multiwell plate. (2) Centrifugation generates a pressure difference *P* across the filter, driving liquid from the apical to the basal chamber. Cells wider than the filter pores must deform to enter the basal chamber (permeate), with mechanically softer cells expected to undergo greater deformation and therefore exhibit higher permeate recovery than stiffer cells. (3) Cell recovery in the permeate is combined with measurements of cell size to obtain a size-corrected proxy for whole-cell stiffness.

Commercial transwell inserts are available with only a limited range of pore sizes. Therefore, we developed modular, 3D-printed inserts that accommodate interchangeable filters (**Figure 2A,B**). Each insert has an inner and outer component which secure a polyester track-etched membrane with cylindrical, monodisperse pores (**Methods**). To evaluate the performance of the inserts, we centrifuged *Escherichia coli*, which has an average cell of width of 1.04±0.01 μm (mean±SD), through filters with pore diameters expected to prevent or permit cell transport. Only 0.0014±0.0035% (mean±SD) of stock cells were recovered in the permeate when 0.4-µm pores were used, indicating that cells did not bypass the filter or pass through pores substantially smaller than the cell width. By contrast 92.5±8.81% of stock cells were recovered in the permeate when the pore diameter was 3.0 µm, which is substantially larger than cell width (**Figure 2C**). These results indicate that nonspecific retention of cells to nonporous regions of the membrane is limited and that most cells can be recovered following centrifugation when deformation is not required.

**Figure 2.**
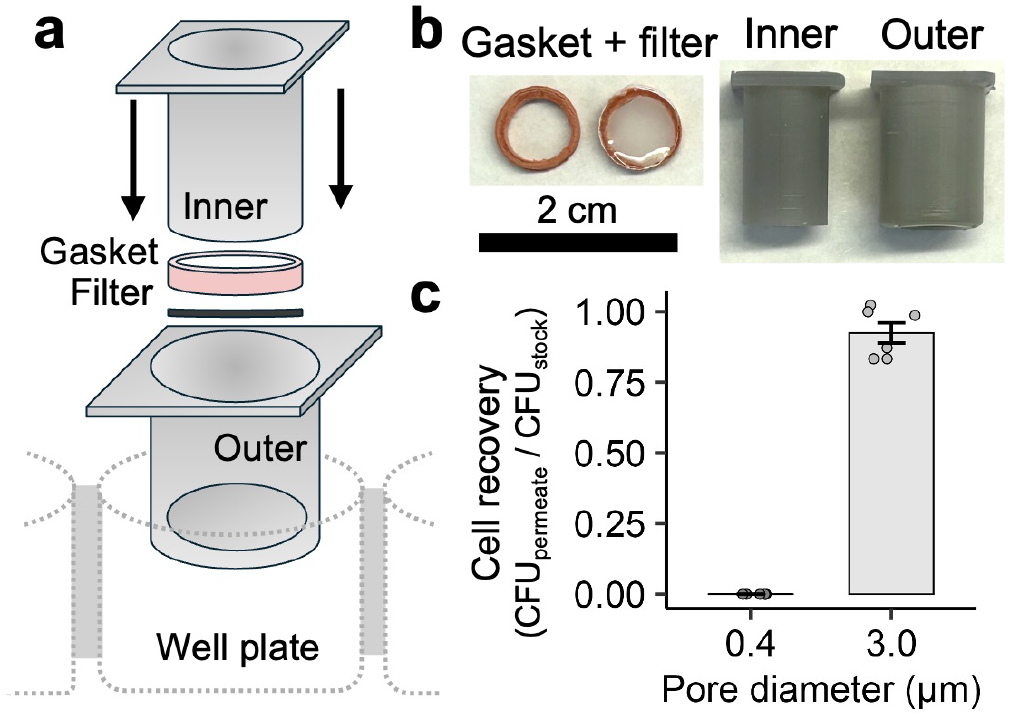
Design and validation of custom transwell inserts for mechanofiltration. (A) Schematic of the 3D-printed transwell insert showing interlocking inner and outer components which secure a polyester track-etched filter and gasket within a multiwell plate. (B) Photograph of the assembled filter and gasket together with the inner and outer insert components. (C) Validation of the insert design using *E. coli*. Essentially no cells were recovered through 0.4-µm pores, which are too small to permit cell transport, but nearly complete recovery was obtained through 3.0-µm pores, which permit cell passage without deformation. These results indicate minimal nonspecific cell retention by the insert and efficient recovery when deformation is not required. Data are shown as mean±SD (*N*=6 biological replicates from two independent experiments).

Cell recovery can be influenced by factors other than stiffness, including filter clogging, loss of cell viability, and variation in cell size. We therefore established experimental and analytical procedures to limit or account for these factors using *E. coli* as a model organism.

#### Filter clogging

Accumulation of cells on the apical surface of the filter can obstruct pores and reduce the transport of subsequently arriving cells. Clogging would decrease permeate recovery and thereby produce an apparent increase in cell stiffness. This effect is minimized by reducing the density of cells in the stock suspension (**Figure 3**). For *E. coli*, stock densities below 10^7^ colony forming units per milliliter (CFU/mL) produced maximal and consistent recovery through filters with 1.0-μm pores. Smaller pores reduced cell transport and increased cell accumulation on the apical surface, requiring lower stock densities to avoid clogging. For example, maximal recovery through 0.8-μm pores was observed only when the stock density was below 10^6^ CFU/mL (**Figure S1**). Thus, the appropriate stock-density range should be established using smaller pore diameters.

**Figure 3.**
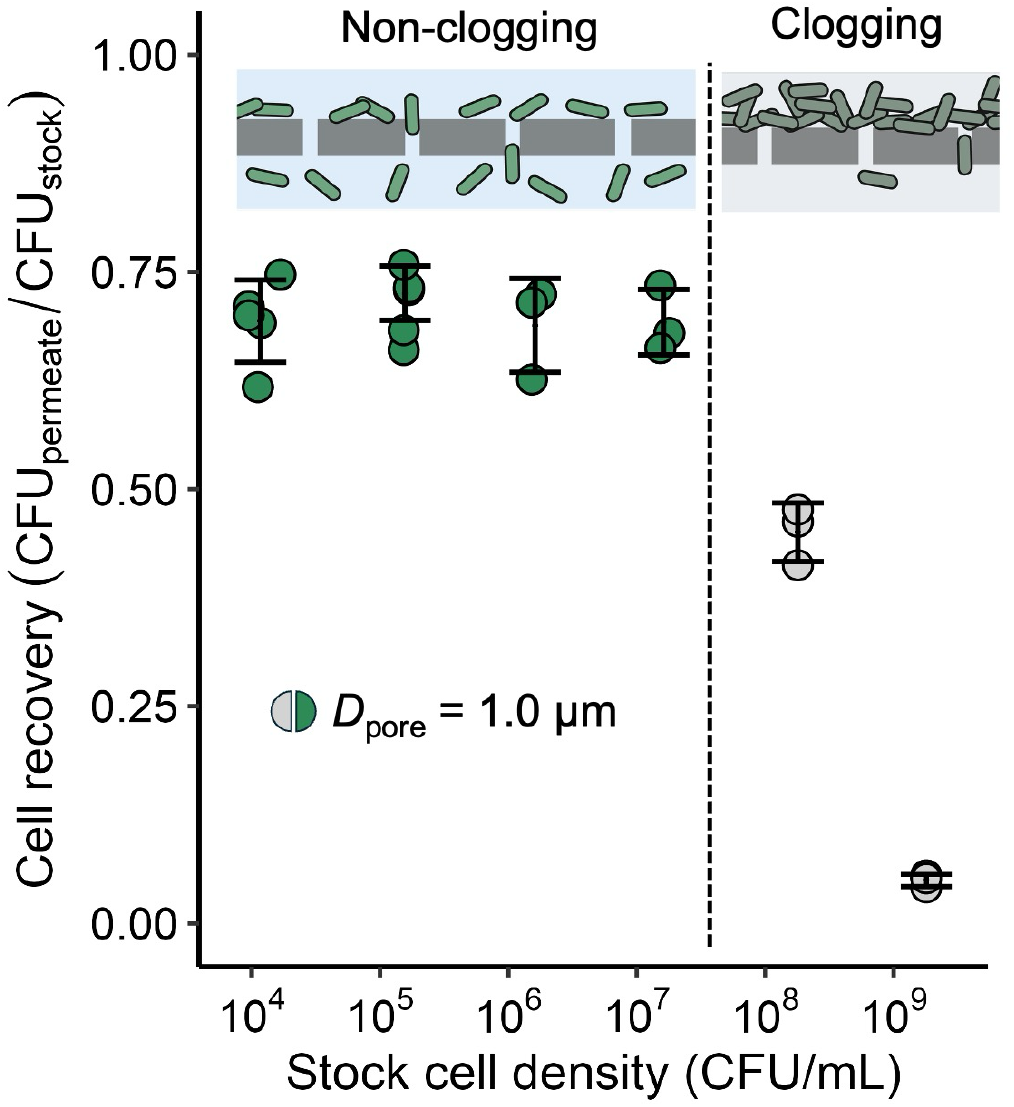
Effect of stock cell density on mechanofiltration recovery. *E. coli* cell recovery following mechanofiltration through 1.0-µm filters is shown as a function of the starting stock cell density. At high cell densities (>10^7^ CFU/mL), accumulation of cells on the apical side of the filter leads to clogging and reduces cell recovery in the permeate (gray). Below this threshold, cell recovery remains constant (green), indicating that clogging is minimized and permeate recovery is independent of stock cell density. Data are shown as mean±SD (*N*≥3 biological replicates from at least two independent experiments).

#### Cell viability and membrane integrity

Passage through a pore may damage the cell envelope or cause lysis when substantial deformation is required. Because the assay quantifies viable cells, deformation-induced loss of viability would reduce permeate recovery and lead to an overestimate of cell stiffness. To identify pore diameters suitable for analysis of *E. coli*, we quantified viable cells recovered from both the permeate and the apical side of the filter. If transport caused extensive cell death or lysis, the combined number of cells recovered from these compartments, CFU_permeate_+CFU_apical_, would be substantially lower than the number of cells present in the initial stock, CFU_stock_.

For filters with pore diameters of 1.0, 0.9, or 0.8 μm, at least ∼75% of the original stock cells were recovered from the apical and permeate compartments (**Figure 4A**). Cells recovered in the permeate retained normal morphology, and little cell debris was visible, consistent with a low incidence of lysis (**Figure 4B**). Thus, some unrecovered cells may still be retained on the insert after washing. When the pore diameter was reduced to 0.7 μm, only 35.5±5.6% of the initial cells were recovered, and cell debris was evident in the permeate (**Figure 4B**). Passage through pores substantially smaller than average cell width can therefore cause appreciable cell lysis.

**Figure 4.**
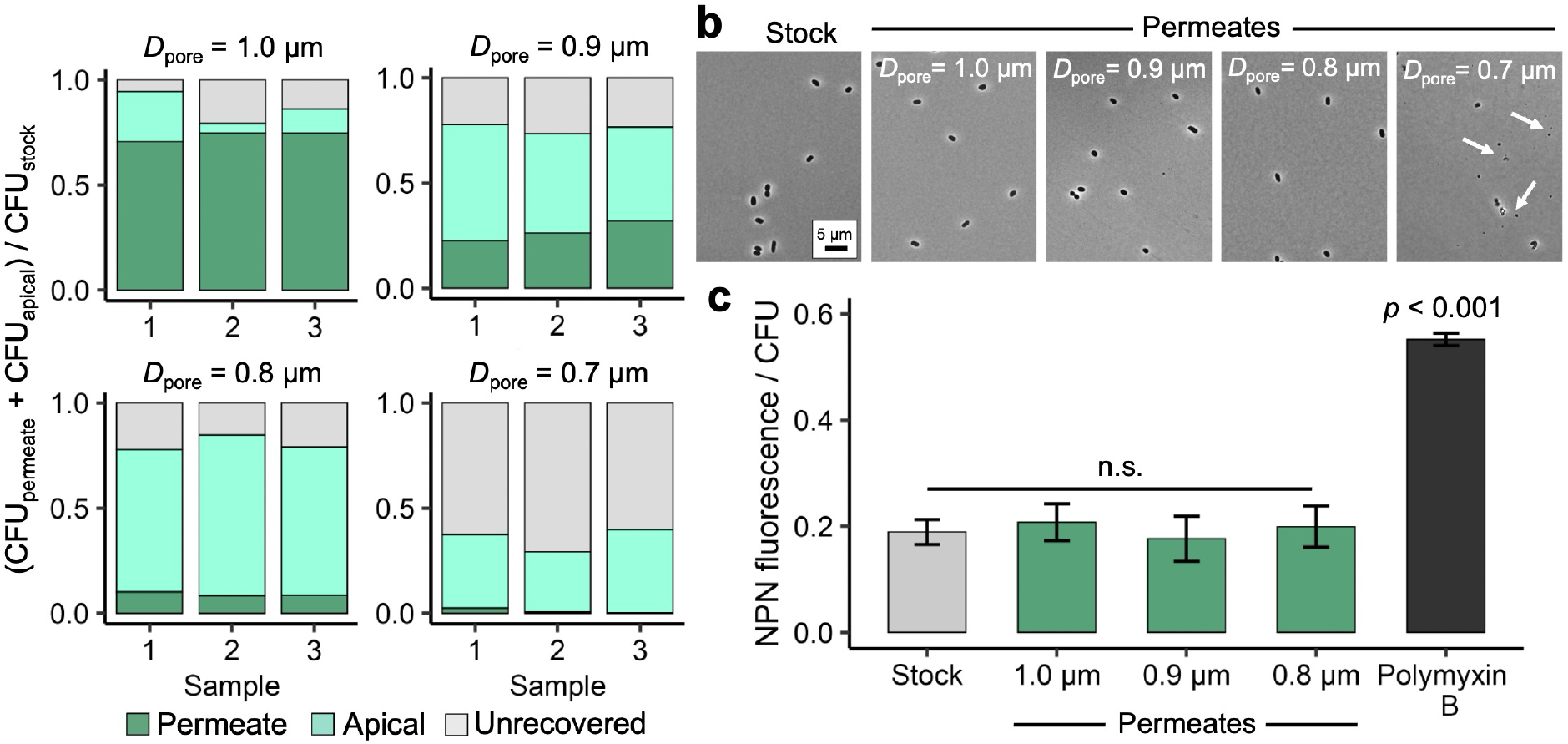
Assessment of cell viability and outer membrane integrity following mechanofiltration. (A) Recovery of viable *E. coli* cells from the permeate and apical chamber following mechanofiltration through filters of the indicated pore diameters. The unrecovered fraction represents cells not recovered from either compartment and is consistent with cell loss during transport. Recovery exceeded ∼75% for filters with pore diameters of 0.8 µm or greater but decreased markedly for 0.7-µm filters, consistent with substantial cell lysis. (B) Representative phase-contrast images of stock cells and cells recovered from permeates. Cells retained normal morphology following passage through 1.0-, 0.9-, and 0.8-µm filters, whereas cell debris (arrows) was evident in permeates collected from 0.7-µm filters. Scale bar, 5 µm. (C) Outer membrane integrity assessed by NPN uptake. NPN fluorescence was normalized to CFU and compared among untreated stock cells, cells recovered from permeates following mechanofiltration, and polymyxin B-treated cells (positive control). Passage through 1.0-, 0.9-, or 0.8-µm filters did not increase NPN uptake relative to stock cells, indicating that mechanofiltration under these conditions did not detectably compromise outer membrane integrity. Data in (A) are from three independent experiments. Data in (C) are shown as mean±SD (*N*=6 biological replicates from two independent experiments). Statistical comparisons were performed using Dunnett’s test versus the stock control; n.s., not significant (*p*>0.05).

We next tested whether passage through the larger pores caused outer-membrane damage without eliminating viability. Outer-membrane permeability was assessed using 1-N-phenylnaphthylamine (NPN), a hydrophobic fluorescent probe whose fluorescence increases when it enters a compromised outer membrane. Polymyxin B-treated cells served as a positive control for membrane permeabilization **(Figure 4C)**. Cells recovered after passage through 1.0-, 0.9-, or 0.8-μm pores did not show increased NPN fluorescence relative to untreated stock cells, indicating that passage through these pores did not detectably compromise outer-membrane integrity **(Figure 4C)**^49,50^.

#### Accounting for variation in cell size

Cell size varies within a clonal population and can also differ across strains or treatment conditions. Smaller cells require less deformation to pass through a pore, and cells narrower than the pore can enter the permeate without deforming. We confirmed that variation in cell size affected transport in mechanofiltration. Relative to the starting stock, the mean width of cells recovered in the permeate decreased by 20 nm for 1.0-μm pores, 40 nm for 0.9-μm pores, and 90 nm for 0.8-μm pores, demonstrating enrichment of narrower cells in the permeate (**Figure 5A**). Permeates nevertheless contained cells whose measured widths exceeded the pore diameter, consistent with deformation during transport. Because cell widths in the starting *E. coli* population were approximately normally distributed^51,52^, we estimated the fraction of stock cells narrower than each pore diameter (**Methods**). These fractions were 32.0% for 1.0-μm pores, 8.5% for 0.9-μm pores, and 1.0% for 0.8-μm pores. In each case, measured recovery exceeded the estimated fraction of cells narrower than the pores, further supporting the conclusion that some cells wider than the pores deformed to enter the permeate (**Figure 5B**).

**Figure 5.**
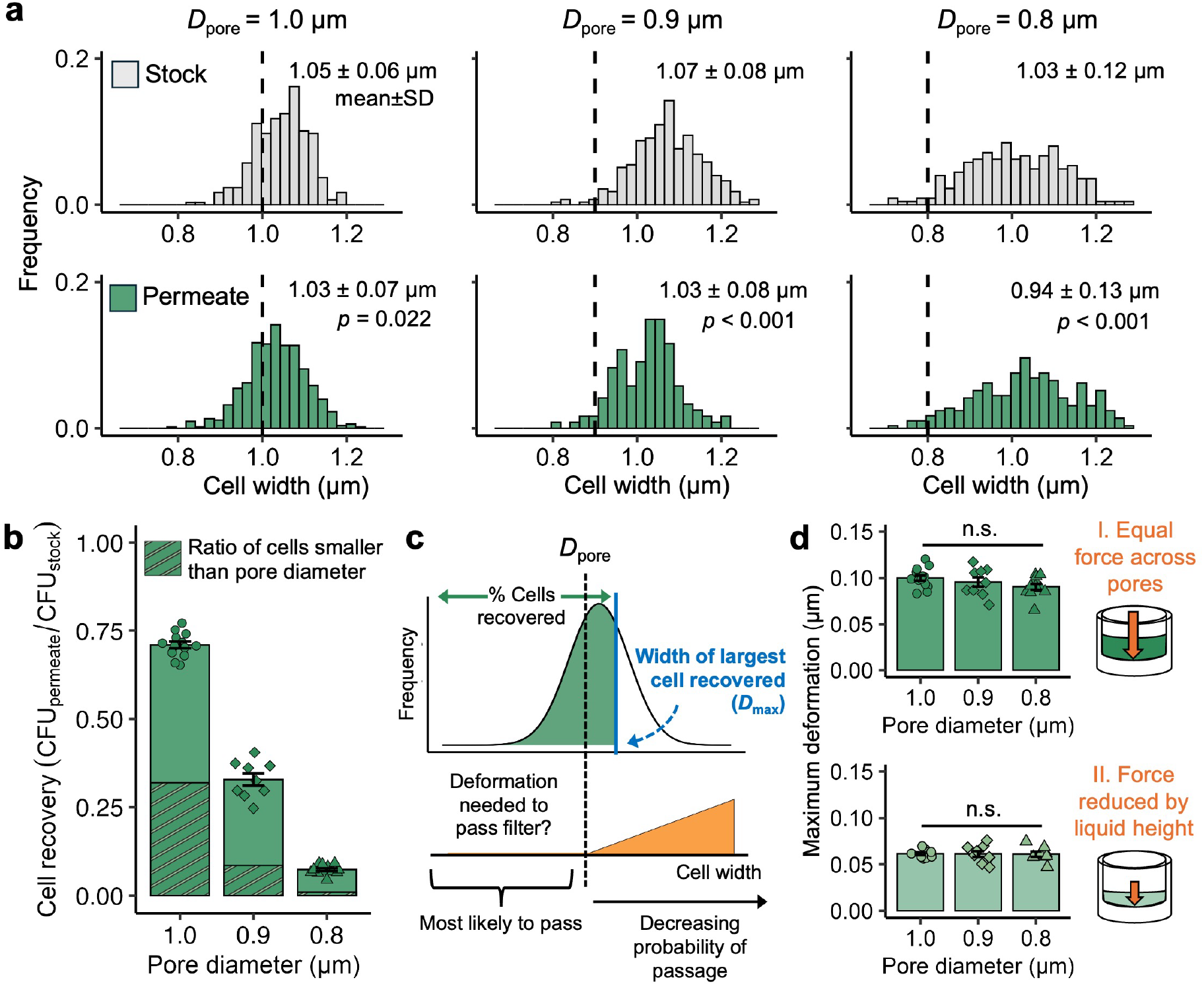
Cell size-corrected maximum deformation provides a pore size-independent measure of bacterial mechanics. (A) Cell-width distributions for *E. coli* in the starting population and among cells recovered in the permeate following mechanofiltration through filters with pore diameters of 1.0, 0.9, and 0.8 µm. Permeates are enriched for smaller cells, although cells wider than the pore diameter (dashed line) are also recovered, indicating deformation during transport. (B) Measured cell recovery exceeds the estimated fraction of cells narrower than the filter pores (hatched region), demonstrating that cells larger than the pore diameter reach the permeate. (C) Schematic illustrating the calculation of maximum deformation (Δ_max_). Assuming that the probability of transport decreases with increasing cell width, the width of the largest cell predicted to reach the permeate (*D*_max_) can be estimated from the measured recovery and the distribution of stock cell widths. (D) Maximum deformation is independent of pore diameter when the same loading conditions are applied (I) but decreases when the applied pressure is reduced by decreasing the height of the liquid column in the apical chamber (II). Data in (A) include >100 cells per condition collected from 3–4 permeates and compared with time-matched stock populations. Statistical comparisons in (A) were performed using Dunnett’s test versus the stock population. Data in (B) represent mean±SD (*N*=9 biological replicates from three independent experiments); hatched regions indicate the estimated fraction of stock cells narrower than the pore diameter. Data in (D) are shown as mean±SD (*N*=9 for standard loading and *N*=6 for reduced loading from at least two independent experiments). Statistical comparisons were performed using one-way ANOVA with Tukey’s post hoc test; n.s., not significant (*p*>0.05).

Cell recovery alone cannot provide a direct measure of stiffness, due to the effect of cell size on transport. Therefore, we defined an inferred metric termed maximum deformation (Δ_max_). This quantity represents the difference between the pore diameter and the estimated width of the largest cell capable of reaching the permeate under a given assay condition. To calculate Δ_max_, we used the measured recovery fraction and the distribution of widths in the stock population to estimate the width of the largest transported cell, *D*_max_. This calculation assumes that the likelihood of transport is ordered primarily by cell width, such that narrower cells pass through more readily than wider cells under the same loading condition (**Figure 5C**). *D*_max_ was obtained from the inverse cumulative normal distribution of stock-cell widths using the measured recovery fraction as the cumulative probability (**Methods**). Maximum deformation was then calculated as *D*_max_ minus the pore diameter. For *E. coli*, Δ_max_ was similar across filters with pore diameters of 1.0, 0.9, and 0.8 μm, with an average of 0.10±0.01 μm (**Figure 5D**). Thus, although raw recovery depended strongly on pore diameter, the size-adjusted maximum-deformation metric was comparatively insensitive to pore size over this range.

To examine this behavior theoretically, we developed an analytical model describing the passage of a deformable cell through a cylindrical pore (**Theory** section in **Supplemental Text**). The model was adapted from the analytical treatment of press-fit components^53^ and approximates the cell as a homogenous solid driven longitudinally through a hollow cylinder with effectively infinite wall thickness (the filter pore). Under these assumptions, the model predicts that maximum deformation depends on the mechanical properties of the cell and filter, including stiffness and Poisson’s ratio (which were maintained across our experiments), and on the applied load. In agreement with our experimental findings, maximum deformation remains independent of pore diameter over the conditions considered.

To evaluate our analytical model, we experimentally altered the loading condition on cells by reducing the volume of stock suspension added to the apical chamber from 200 to 100 µL. This reduction decreases the height of the liquid column and therefore lowers the pressure generated across the filter during centrifugation (**Supplemental Equation 1**). Cell recovery decreased at the lower volume, consistent with reduced transport of wider cells into the permeate (**Figure S2**). The corresponding maximum deformation decreased by 36% (**Figure 5E**). This response supports the expected dependence of cell deformation on the applied loading condition and is broadly consistent with the scaling predicted by the analytical model.

### Maximum Deformation Reports Changes in Bacteria Cell Stiffness

We next evaluated whether mechanofiltration could detect established changes in *E. coli* cell stiffness. We examined two genetic perturbations (Δ*ompA* and Δ*lpp*) and two chemical perturbations (ethylenediaminetetraacetic acid (EDTA) and A22), each of which has been shown to reduce *E. coli* cell stiffness^39,41,54–58^. As reported previously, all four perturbations increased average cell width relative to unperturbed wild-type cells (**Figure S3**)^58^. Raw cell recovery increased for Δ*ompA*, EDTA-treated, and A22-treated cells, consistent with greater deformation during transport through the filter (**Figure 6A**). In contrast, Δ*lpp* cells exhibited lower cell recovery than wild type despite being mechanically softer. These cells also had substantially larger widths than all other groups examined (1.19±0.07 µm), again illustrating that cell recovery is strongly influenced by cell size and cannot be interpreted as a direct measure of cell stiffness.

**Figure 6.**
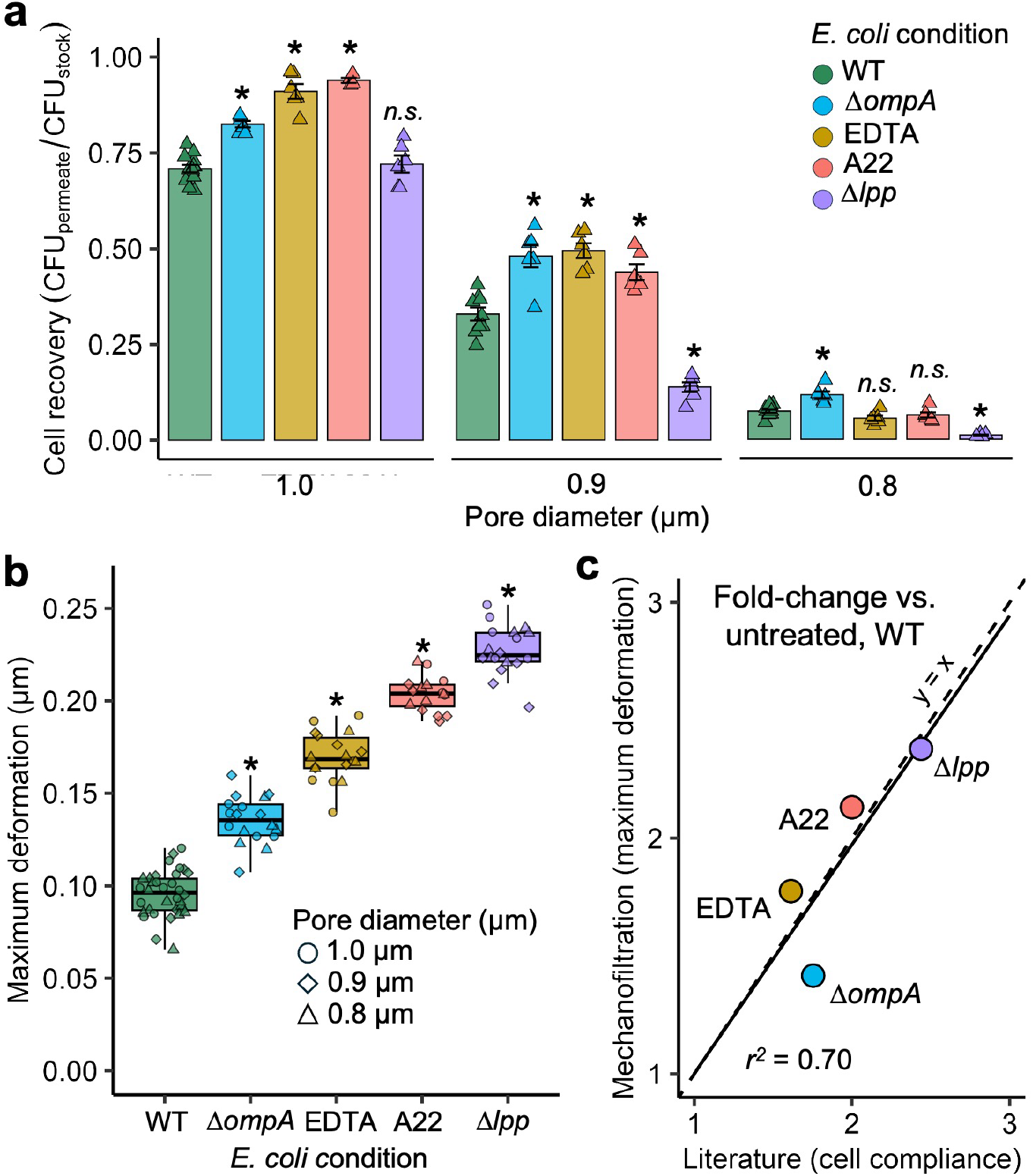
Mechanofiltration detects established changes in *E. coli* cell mechanics. (A) Cell recovery following mechanofiltration for untreated wild-type (WT) *E. coli* and cells with reduced stiffness resulting from genetic (*ΔompA, Δlpp*) or chemical (EDTA, A22) perturbations. Raw cell recovery varies among conditions and is influenced by differences in cell width. (B) Maximum deformation (Δ_max_) calculated from cell recovery and stock cell-width distributions. All four perturbations significantly increased Δ_max_ relative to untreated WT cells, consistent with reduced whole-cell stiffness. Symbols indicate measurements obtained using different filter pore diameters. (C) Fold-changes in Δ_max_ relative to untreated WT cells compared with published measurements of cell compliance obtained using AFM, cell-bending assays, or osmotic shock experiments. Published data are expressed as fold changes in cell compliance (the inverse of stiffness). Mechanofiltration reproduces the relative magnitude of stiffness changes reported using established methods. Data in (A) and (B) are shown as mean±SD (WT*, N*=9; all other conditions, *N*=6 biological replicates from at least two independent experiments). Data in (C) represent mean values from this study compared with published (ref. 39, 58). Statistical comparisons in (A) and (B) were performed using Dunnett’s test versus WT. *: *p*<0.01. n.s., not significant (*p*>0.05).

To account for the effects of cell size, we calculated maximum deformation (Δ_max_) for each condition. As observed above, Δ_max_ was largely independent of pore diameter, with only a small but statistically significant difference among filters for Δ*lpp* mutants (**Figure S4**). All four perturbations increased Δ_max_ relative to unperturbed wild-type cells (*p<*0.001 for each comparison) (**Figure 6B**), consistent with reduced cell stiffness.

To benchmark mechanofiltration against established approaches, we compared fold changes in Δ_max_ with published measurements of cell stiffness for the same perturbations obtained using AFM, cell-bending assays, or osmotic shock experiments (**Table S1**)^39,55,58^. Fold changes measured by mechanofiltration closely tracked those reported previously (Pearson’s *r*=0.84; **Figure 6C**). Although mechanofiltration does not directly measure material properties such as Young’s modulus, it reproduced the relative magnitude of stiffness changes across diverse genetic and chemical perturbations while utilizing substantially simpler instrumentation.

### Mechanofiltration Detects Stiffness Differences Across Bacterial Species

We next asked whether mechanofiltration could distinguish differences in cell stiffness across bacteria with diverse morphologies and cell envelope architectures. To test the generality of the assay, we measured Δ_max_ for the Gram-negative species *Vibrio cholerae* and the Gram-positive species *Bacillus subtilis* and *Staphylococcus aureus* (**Figure S5**). Previous studies, including work from our laboratories, have shown that *V. cholerae* is substantially softer than *E. coli*^41,58^, *B. subtilis* is substantially stiffer^37,40^, and *S. aureus* exhibits comparable mechanics to *E. coli*^41,59^.

Because clogging behavior may differ among species, we first established stock densities that produced consistent permeate recovery. Clogging was avoided at concentrations below 10^4^ CFU/mL for *V. cholerae* and *B. subtilis*, and below 10^5^ CFU/mL for *S. aureus* (**Figure S6**). These differences suggest that species-specific characteristics, including cell shape, aspect ratio, surface chemistry, and cell-cell interactions, influence filter clogging during mechanofiltration.

Using these optimized conditions, we measured cell recovery with at least two pore diameters for each species to calculate Δ_max_ (**Figure 7, Figure S7**). Mechanofiltration reproduced the relative differences in stiffness reported previously. *V. cholerae*, which has previously been shown to be the softest of these organisms^41,58^, exhibited the largest maximum deformation (0.16±0.02 μm; 0.8-, 0.7-, and 0.6-µm pores). The characteristic curved morphology of *V. cholerae* did not appear to influence transport, as the distribution of cell curvatures in the permeate was similar to that of the starting population (**Figure S8**). By contrast, *B. subtilis*, previously reported to be substantially stiffer than *E. coli*^37,40^, exhibited no detectable deformation (Δ_max_=0.00±0.02 μm; 0.8-μm and 0.7-μm pores). Consistent with this result, the average width of cells recovered in the permeate was less than the pore diameter (**Figure S9**), indicating that transport was largely restricted to cells that could pass through the filter without measurable deformation. Finally, Δ_max_ for *S. aureus* (0.10±0.01 μm; 1.0-μm and 0.9-μm pores) was indistinguishable from that of *E. coli* (*p*=0.12). Although *S. aureus* is a Gram-positive organism and may thus be expected to have higher cell stiffness due to a thicker cell wall, we previously observed that *S. aureus* deformation was slightly greater than *E. coli* under the same loading conditions within our microfluidic device^41^. This observation is consistent with our results from mechanofiltration, as well as AFM studies showing marginal differences in cell mechanics between *S. aureus* and *E. coli*^59^.

**Figure 7.**
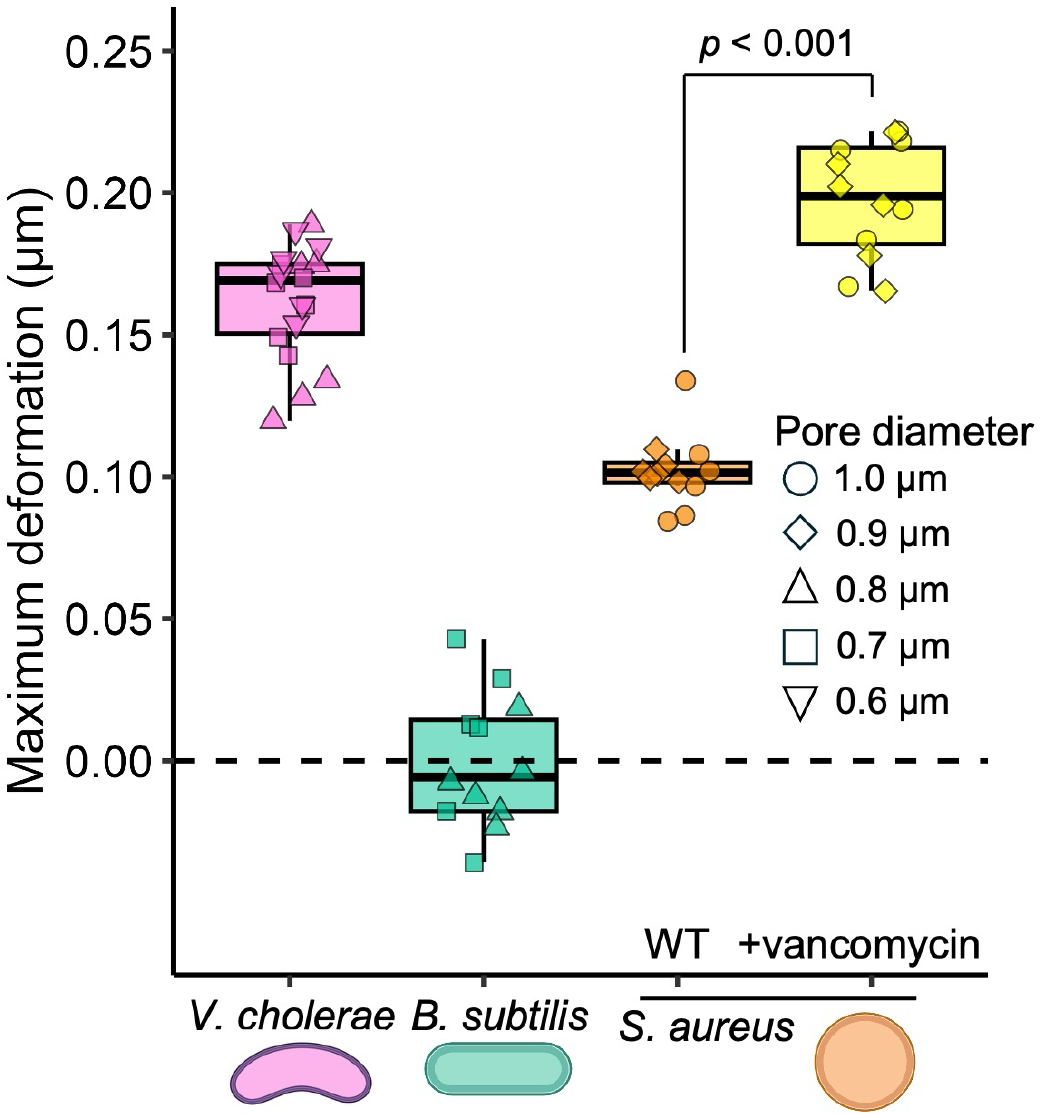
Mechanofiltration detects differences in bacterial mechanics across species. Maximum deformation (Δ_max_) measured by mechanofiltration for wild-type *V. cholerae*, *B. subtilis*, and *S. aureus*. Multiple filter pore diameters (1.0, 0.9, 0.8, 0.7, and/or 0.6 µm, as indicated) were used to calculate Δ_max_ for each species. Consistent with previous reports, *V. cholerae* exhibited the largest maximum deformation, *B. subtilis* showed no detectable deformation, and *S. aureus* exhibited deformation comparable to *E. coli*. Treatment of *S. aureus* with 40 µg/mL vancomycin for 1 h significantly increased Δ_max_, consistent with reduced whole-cell stiffness following inhibition of cell-wall crosslinking. Data are shown as mean±SD (*N*=6 biological replicates per filter from two independent experiments). Statistical comparisons between untreated and vancomycin-treated *S. aureus* were performed using a two-tailed Student’s t-test.

To determine whether mechanofiltration could also detect changes in cell stiffness within bacterial species other than *E. coli*, we treated *S. aureus* with vancomycin, which inhibits peptidoglycan crosslinking^60,61^. Although the biomechanical consequences of vancomycin treatment have not been directly measured in *S. aureus*, genetic disruption of cell-wall crosslinking has previously been shown to reduce cell-wall stiffness^62^. Cells were treated with vancomycin (40 μg/mL, 10X MIC) for 1 h under conditions that did not affect cell viability^61^. Vancomycin increased the average cell width from 1.08±0.06 μm to 1.15±0.08 μm (*p*<0.001), consistent with cell-wall softening and expansion under turgor pressure (**Figure S10**). Correspondingly, mechanofiltration detected a twofold increase in maximum deformation, from 0.10±0.01 µm to 0.20±0.02 μm following vancomycin treatment (**Figure 7**). Together, these results demonstrate that mechanofiltration can distinguish both interspecies and intraspecies differences in bacterial cell stiffness.

## DISCUSSION

Mechanofiltration provides a simple, inexpensive approach for assessing bacterial biomechanics using standard laboratory equipment and readily fabricated transwell inserts. Unlike existing methods that rely on AFM, microfluidic devices, or specialized instrumentation, mechanofiltration can be implemented with common microbiological workflows while maintaining sensitivity to biologically meaningful differences in whole-cell mechanics. Across a range of genetic, chemical, and species-level perturbations, the assay reproduced trends in bacterial stiffness previously established using more sophisticated approaches, supporting its utility as a rapid screening tool for bacterial biomechanics.

The principal output of mechanofiltration is the maximum deformation metric, Δ_max_, which provides a size-corrected measure of whole-cell deformability rather than a direct estimate of material properties such as Young’s modulus. More detailed mechanical information can be obtained using techniques such as AFM together with mechanical modeling. In contrast, mechanofiltration prioritizes accessibility and throughput. However, the assay also has practical limitations. Measurements require preliminary optimization to avoid filter clogging when new species are examined, and our current implementation relies on colony counting to quantify permeate recovery, limiting throughput. In addition, our experiments were performed with stationary-phase cells to prevent proliferation during the assay, whereas many published biomechanical measurements have been obtained during exponential growth^39,41,55,58^. Because cell-envelope composition changes with growth phase^63–65^, these differences may contribute to quantitative differences between mechanofiltration and previous measurements^39,41,55,58^. Future implementations incorporating automated cell quantification^66–69^ should further improve assay throughput.

The simplicity and scalability of mechanofiltration make it well suited for screening large collections of bacterial strains and experimental conditions. Such studies could identify new genetic and environmental determinants of bacterial mechanics, which may serve as antimicrobial targets or indicators of antibiotic susceptibility. We anticipate that mechanofiltration will provide an accessible platform for expanding the study of bacterial biomechanics beyond laboratories with specialized instrumentation. Interestingly, we and others have found that sublethal exposure to antibiotics can reduce bacterial cell stiffness, including A22 treatment in *E. coli* and vancomycin treatment in *S. aureus*^39–41,70^. Antibiotic-induced changes in bacterial mechanics may influence further drug susceptibility and colonization by enabling softer cells to traverse submicron cavities within tissues and materials, where they may be protected from host immunity, sterilization procedures, and antibiotic exposure^44^. More broadly, mechanofiltration provides an accessible platform for identifying conditions that alter bacterial biomechanics and for prioritizing strains and perturbations for detailed investigation.

## METHODS AND MATERIALS

### Bacterial Strains and Growth

Bacterial strains were recovered from glycerol stocks and grown overnight (14-18 h) at 37 °C with shaking (220 rpm). *E. coli* MC4100*, V. cholerae* C6707, and *B. subtilis* 168 were cultured in lysogeny broth (LB; Sigma-Aldrich #L2542). *S. aureus* SA25923 was cultured in Tryptic Soy Broth (TSB; Millipore Sigma, #146317). LB cultures of *E. coli* knockout mutants were supplemented with 50 μg/mL kanamycin (Millipore Sigma, #K1377). The *E. coli* knockout strains have been described previously^58^.

To alter bacterial mechanics, wild-type *E. coli* cultures were treated immediately before mechanofiltration with either 10 mM EDTA (Research Products International, #E14000) or 10 μg/mL A22 (Sigma Aldrich, #SML0471) for 10 min at room temperature. *S. aureus* suspensions were treated with 40 μg/mL vancomycin (Millipore Sigma, #V2002) for 1 h at 37 °C. Treated cultures were subsequently diluted by 10^−6^-fold to reduce the antibiotic concentration to subinhibitory levels while simultaneously achieving the cell density required for mechanofiltration.

### Insert Design and Fabrication

Custom transwell inserts were designed for 48-well plates (Corning Costar, #3548) using Fusion360 (Autodesk, San Francisco, CA, USA). Each insert consists of interlocking inner and outer cylindrical components that secure a porous filter. The 48-well inserts measure 13.25 mm in length, with inner diameters of 6.5 mm (inner component) and 8.8 mm (outer component). The outer component includes a 2-mm-wide circumferential ledge that supports the filter. Both components terminate in square caps (16.5×16.5 mm) that rest on the plate to suspend the insert within each well. Inserts were fabricated using a Form 3 stereolithography printer (Formlabs, Somerville, MA USA) with Tough 1500 resin.

Larger transwell inserts were fabricated for selected experiments in 24-well plates. For these inserts, the inner and outer diameters were increased to 9.7 mm and 13.6 mm, respectively, while the cap size was increased to 16.6 mm. All remaining dimensions were unchanged.

Polyester track-etched filters (Sterlitech Corporation, Auburn, WA, USA) with pore diameters of 0.4, 0.6, 0.7, 0.8, 0.9, 1.0, or 3.0 μm were used for mechanofiltration (catalog numbers PET0413100, PET1300080, PET1300081, PET1300069, PET1300070, and PET1013100, respectively). For 48-well inserts, 13-mm filters were trimmed using metal dies to fit the insert dimensions. Filters were secured to custom rubber gaskets (William H. Harvey Co., #020500) using clear nail polish (L.A. Colors, #NP195). Transwells were assembled by placing the filter and gasket within the outer component and locking them in place with the inner component. Inserts and gaskets were reused following cleaning with 10% bleach and water, whereas filters were discarded after a single use.

### Mechanofiltration Assay

Overnight cultures were resuspended in fresh growth medium and serially diluted to the desired cell densities. Average cell width was determined from representative stock suspensions by imaging cells on agarose pads (see below). Stock suspensions were then added to the apical chamber of assembled transwell inserts, and wells were sealed with an adhesive plate cover (Bio-Rad, # MSB-1001B). Standard experiments in 48-well plates used 200 μL of bacterial suspension. To examine the effect of the applied pressure on cell recovery, the suspension volume was reduced to 100 μL in selected experiments. Plates were centrifuged in an Eppendorf 5810R centrifuge equipped with MTP buckets at 4000 rpm (3,220*g*) for 15 min for *E. coli*, *V. cholerae*, and *B. subtilis*. Preliminary experiments showed that these centrifugation conditions reduced the viability of *S. aureus* (**Figure S11**). Therefore, *S. aureus* samples were centrifuged for 5 min, and a no-filter control was included in all experiments by centrifuging stock suspensions directly in 48-well plates alongside transwell samples.

Following centrifugation, transwell inserts were removed, permeates were resuspended, and bacterial suspensions were plated on agar for colony enumeration. Samples were serially diluted in fresh growth medium as needed to obtain approximately 30-100 colonies per plate after incubation for 12–24 h at 30 °C. At least two replicate plates were prepared for each sample, and colony counts were averaged. Cell recovery was calculated as the ratio of bacterial concentration in the permeate to that in the starting stock suspension. For *S. aureus*, no filter controls were used in place of the starting stock to calculate cell recovery. To recover cells remaining in the apical chamber, inserts were disassembled, submerged in 5 mL of fresh growth medium, and sonicated for four 30 s intervals (2 min total) in a bath sonicator (Emerson Branson Model 2800). The resulting suspensions were plated for colony enumeration as described above.

Maximum deformation (Δ_max_) was calculated from the measured cell recovery and the distribution of cell widths in the starting population. The width of the largest cell predicted to enter the permeate was estimated using the inverse cumulative normal distribution function, where the measured mean and standard deviation of stock cell widths defined the normal distribution and the measured cell recovery served as the cumulative probability. Maximum deformation was then calculated by subtracting the filter pore diameter from *D*_max_.

To estimate the fraction of cells with widths smaller than the filter pore diameter, the cumulative normal distribution of stock cell widths was evaluated at the pore diameter using the measured mean and standard deviation of the stock population.

### Cell Imaging and Morphometric Analysis

To determine cell dimensions, stock, permeate, and apical cell suspensions were concentrated 100- to 1000-fold in growth medium to obtain sufficient cell densities for imaging (>100 cells per field of view). Aliquots (1 μL) were deposited onto 1.5% agarose pads, covered with glass coverslips, and imaged immediately. Phase-contrast images were acquired using an Olympus IX83 inverted microscope (Evident Scientific, Waltham, MA, USA) equipped with a 100X objective (Evident Scientific, Waltham MA, USA, #UPLXAPO100X0). Cell dimensions were quantified using ImageJ (v. 1.53k, 2021) with the MicrobeJ plugin (v. 5.13p, 2024)^71^. Reported cell widths correspond to the maximum cell width determined using the MicrobeJ “fit to shape” function.

### Outer Membrane Permeability Assay

Outer membrane integrity following mechanofiltration was assessed using uptake of the fluorescent probe 1-*N*-phenylnaphthylamine (NPN)^49,50^. Mechanofiltration was performed using 1.0-, 0.9-, and 0.8-μm filters in 24-well plates with 400 μL of bacterial suspension to increase permeate cell numbers for fluorescence detection. Following mechanofiltration, stock and permeate samples were collected, washed once with 5 mM HEPES buffer (Gibco, #15630080), and resuspended in half of the original volume. Aliquots (95 μL) were transferred to black, clear-bottom 96-well plates (Corning Costar, #3904), followed by the addition of 5 μL of 5 mM NPN (Sigma-Aldrich, #104043) prepared in acetone. Plates were incubated for 30 min at room temperature in the dark before fluorescence was measured (excitation/emission: 350/420 nm) using a SpectraMax iD5 microplate reader (Molecular Devices, San Jose, CA, USA).

Background fluorescence, determined from wells containing HEPES buffer and NPN alone, was subtracted from all measurements. Fluorescence values were normalized to colony forming units (CFU) per well, which were determined by plating serial dilutions of the remaining sample volume on LB agar. For positive controls, aliquots of stock cultures were treated with 10 μg/mL polymyxin B (Sigma-Aldrich, #P1004) for 15 min at room temperature, washed, and resuspended in HEPES buffer as described above. Cells were then incubated with NPN in the continued presence of 10 μg/mL polymyxin B for 30 min before fluorescence measurements were collected.

### Statistical Analysis

Statistical analyses were performed in RStudio (v. 2024.12.0+467) using a significance threshold of α = 0.05. Specific statistical tests are described in the corresponding figure legends.

## Data Availability

Source data are provided with the paper.

## Acknowledgements

We thank the staff of the Jacobs Hall Makerspace at the University of California, Berkeley, for assistance with 3D printing. We also thank Peter Turnbaugh for providing *B. subtilis* 168 and *S. aureus* SA25923. This work was supported by the Center for Cellular Construction, a National Science Foundation (NSF) Science and Technology Center under Cooperative Agreement DBI-1548297, and by an NSF BRITE Fellowship (Award 2342239) awarded to C.J.H.. K.G.D. was supported by Ruth L. Kirschstein National Research Service Award Postdoctoral Fellowship F32AI194695. G.J. was supported by an NSF Graduate Research Fellowship. Figures were created in part with Biorender.com.

## Author contributions

K.G.D. and C.J.H. led study conceptualization with assistance from K.C.H. and J.N.E. K.G.D. performed experiments, analyzed and curated data, performed statistical analyses, and wrote the manuscript. J.W.H assisted with data collection and analysis. J.G. designed and fabricated transwell inserts. K.C.H. and J.N.E. consulted on research design and data interpretation and edited the manuscript. C.J.H. designed and directed research and edited the manuscript.

