## Supplemental Figures 1-12, Table S1, and and Text for "Mechanofiltration Enables High-Throughput Measurements of Bacterial Cell Mechanics"

**Supplemental Figures S1-212**

**Supplemental Table S1**

**Supplemental Text – Theory**

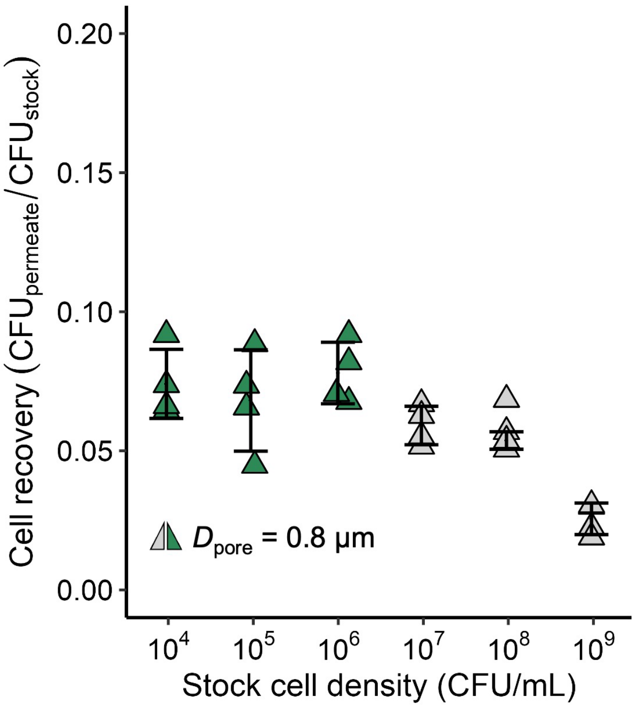


**Figure S1.** *E. coli* cell recovery following mechanofiltration through 0.8-µm filters is shown as a function of the starting stock cell density. At high cell densities (>$10^6$ CFU/mL), accumulation of cells on the apical surface of the membrane reduces permeate recovery because of clogging (gray). Below this threshold, cell recovery remains constant (green), indicating that clogging is minimized and permeate recovery is independent of stock cell density. Data are shown as mean±SD (*N*≥3 biological replicates from at least two independent experiments).

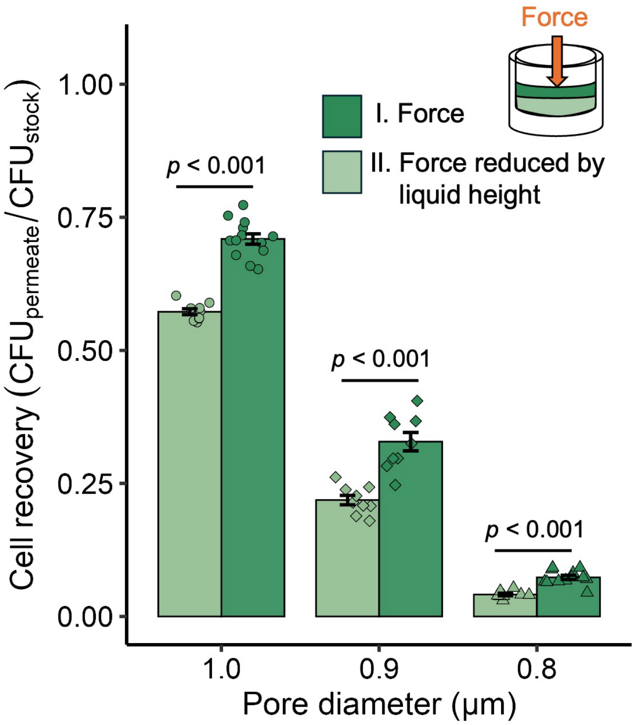


**Figure S2.** Measured cell recovery is dependent on applied force. Standard loading conditions in mechanofiltration (I) are reduced by decreasing the volume of bacterial suspension added to the apical well (II). Reducing force decreases cell recovery, consistent with reduced deformation and transport of larger cells. Data are shown as mean±SD ($N$=9 for standard loading and $N$=6 for reduced loading from at least two independent experiments). Statistical comparisons were performed using Student's t-test.

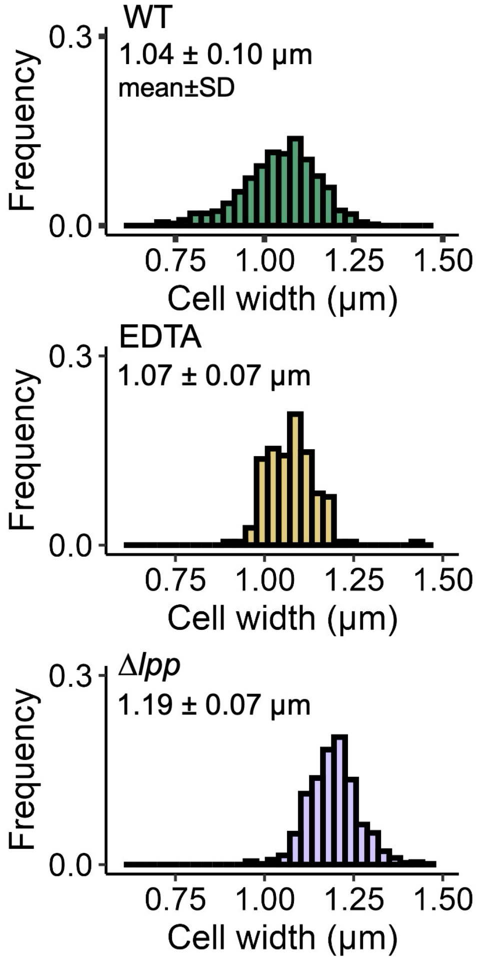


**Figure S3.** Cell-width distributions for stocks of untreated, wild-type (WT) E. coli and E. coli following perturbations to the cell-envelope. Data include >100 cells per condition.

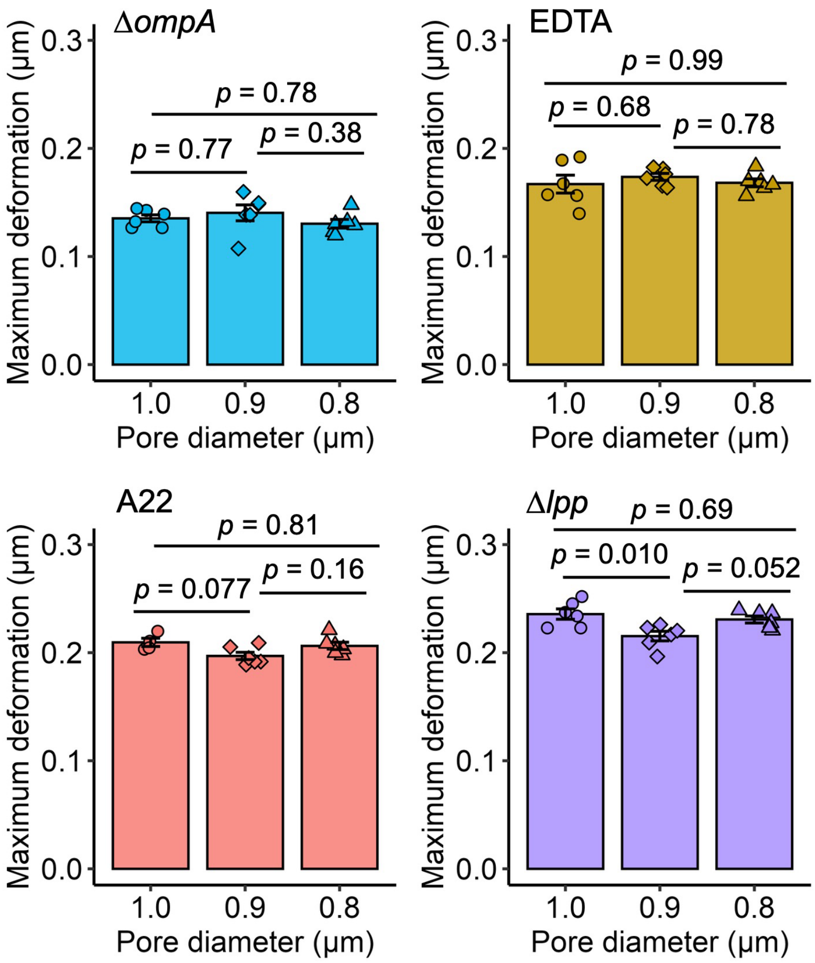


**Figure S4.** Maximum deformation was largely independent of pore diameter for *E. coli* conditions that reduce cell stiffness, with only a small but statistically significant difference among filters for Δ*lpp* mutants. All four perturbations increased maximum deformation relative to unperturbed wild-type cells. Data are shown as mean±SD (*N*=6 biological replicates from at least two independent experiments for each condition). Statistical comparisons were performed using one-way ANOVA with Tukey's post hoc test.

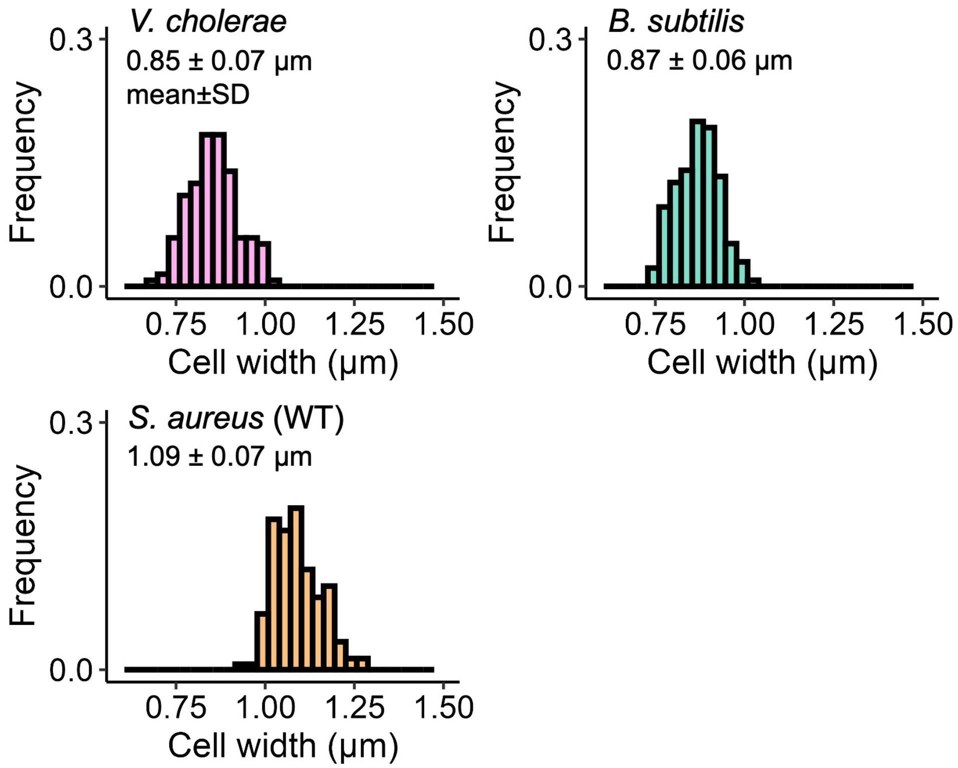


**Figure S5.** Cell-width distributions for *V. cholerae*, *B. subtilis*, and *S. aureus* stocks. Data include >100 cells per condition.

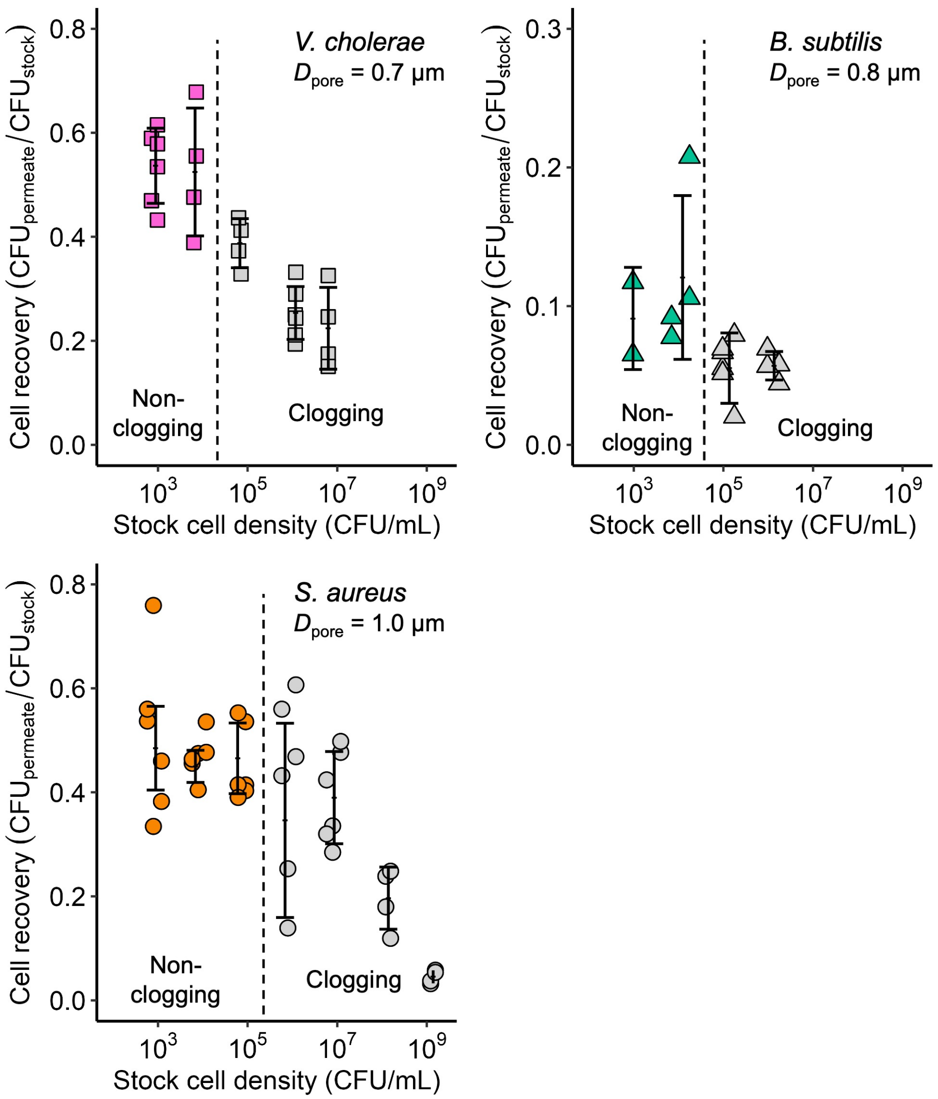


**Figure S6.** Measured cell recovery for *V. cholerae*, *B. subtilis*, and *S. aureus* is shown as a function of the starting stock cell density. At high cell densities (>$10^4$ CFU/mL *V. cholerae* and *B. subtilis*; >$10^5$ CFU/mL *S. aureus*), accumulation of cells on the apical surface reduces permeate recovery because of filter clogging (gray). Below this threshold, cell recovery remains constant (colored), indicating that clogging is minimized and permeate recovery is independent of stock cell density. Data are shown as mean±SD ($N \geq 3$ biological replicates from at least two independent experiments).

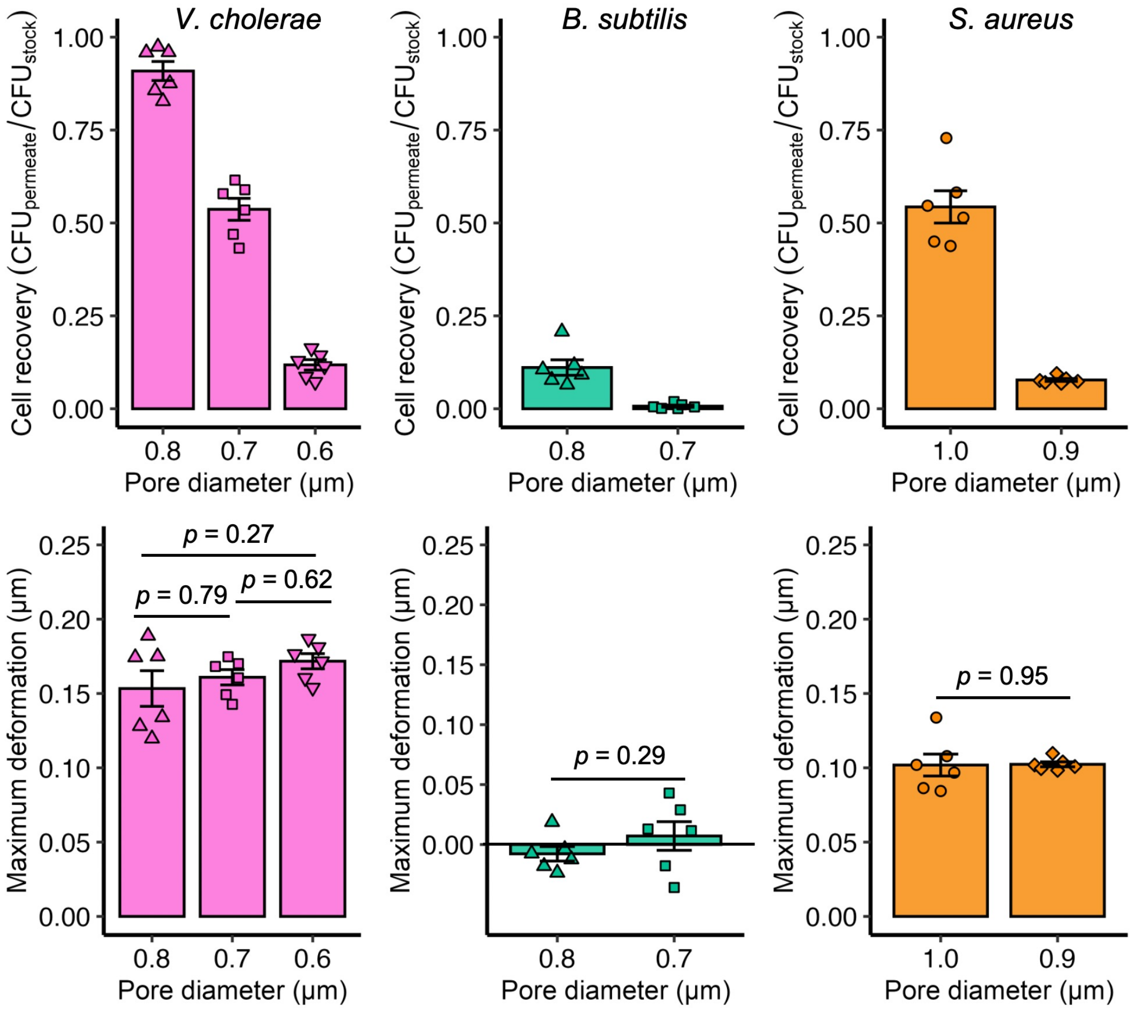


**Figure S7.** Measured cell recovery for *V. cholerae*, *B. subtilis*, and *S. aureus* through filters with specified pore diameters are shown (top) and corresponding maximum deformations (bottom row). Data are shown as mean±SD (*N*=6 biological replicates from at least two independent experiments. Statistical comparisons were performed using one-way ANOVA with Tukey's post hoc test for 3-group comparisons and Student's t-test for 2-group comparisons).

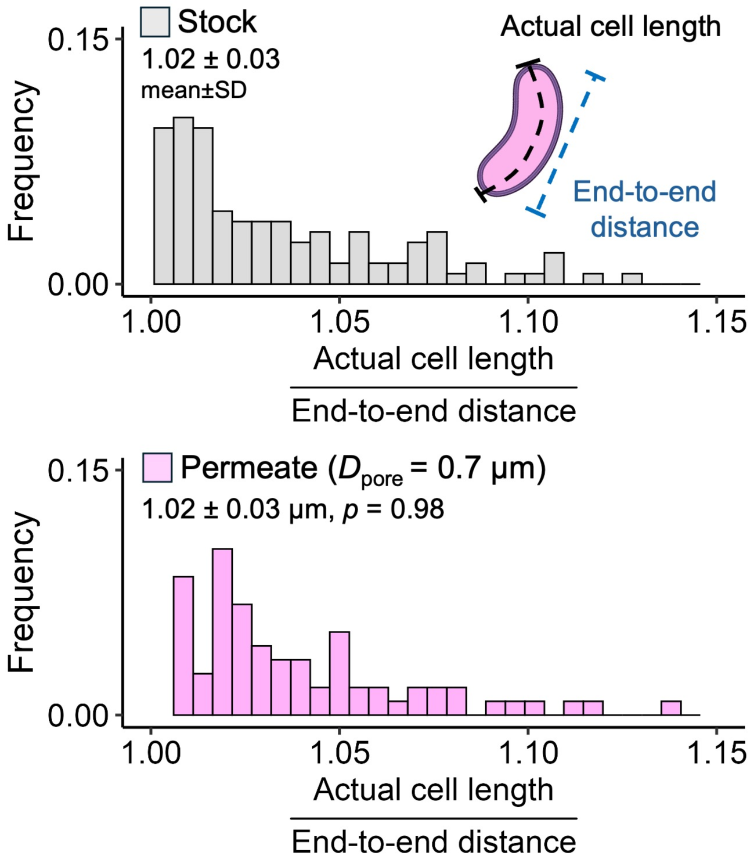


**Figure S8.** Cell curvature is compared for *V. cholerae* before and after mechanofiltration through 0.7-µm filters. Cells in permeates have similar curvature to cells in stocks, assessed by taking the ratio of actual cell length to end-to-end distance. Data include >100 cells per condition collected from 3 permeates and compared with time-matched stock populations. Statistical comparisons in were performed using Dunnett's test versus the stock population.

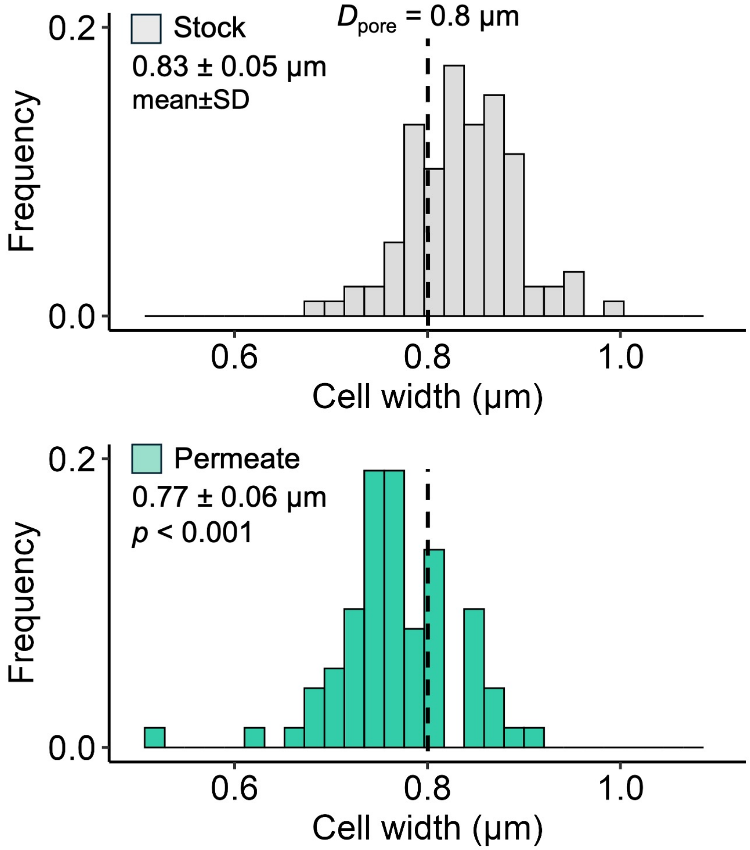


**Figure S9.** Cell-width distributions for *B. subtilis* in the starting population and among cells recovered in the permeate following mechanofiltration through 0.8-μm filters. Permeates primarily include cells that are smaller than pore diameter (dashed line). $N$=98 cells for stock and $N$=73 cells for permeate, obtained from 4 different permeates. Statistical comparisons in were performed using Dunnett's test versus the stock population.

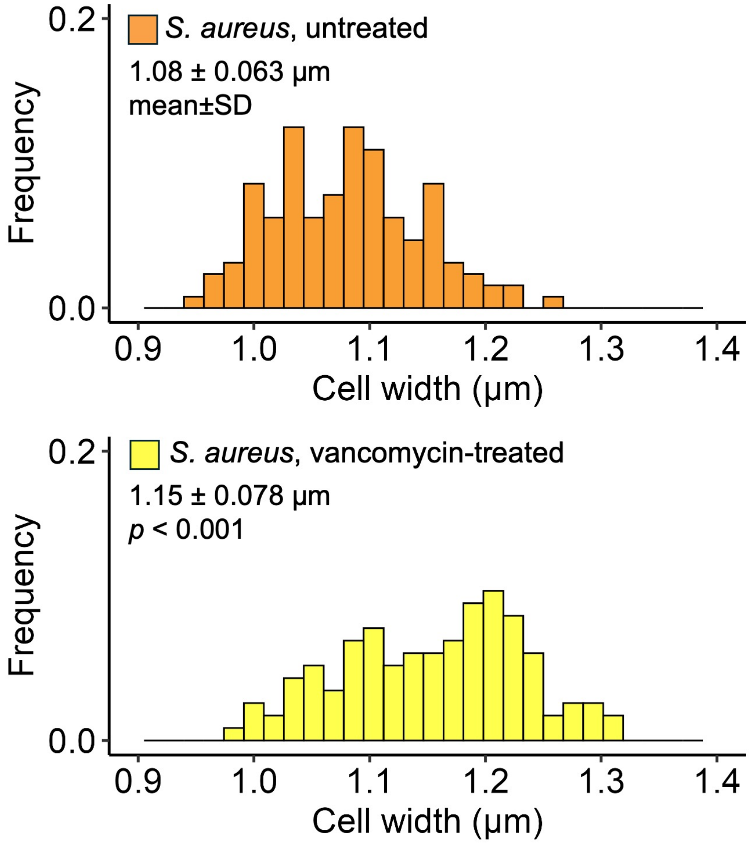


**Figure S10.** Cell-width distributions for *S. aureus* following exposure to vancomycin. Width is compared to time-matched, untreated cells. Data include >100 cells. Statistical comparisons in were performed using Dunnett's test versus untreated group.

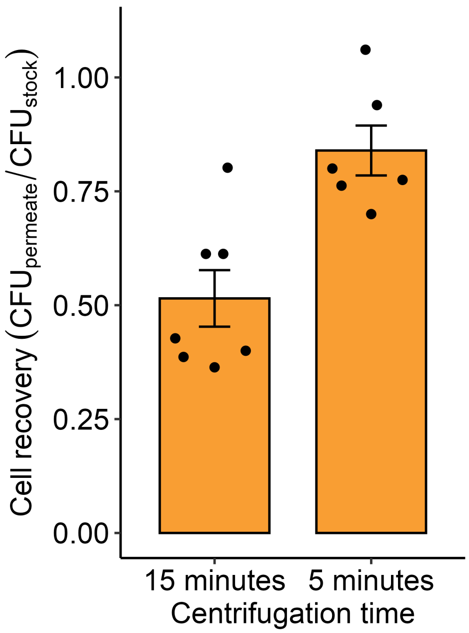


**Figure S11.** Cell recovery of untreated, wild-type *S. aureus* is shown when cells are centrifuged for 5 or 15 minutes without filters. Low recovery at 15 minutes suggests cell death. Cell viability is not affected when time is decreased to 5 minutes. Data are shown as mean±SD (*N*=6 biological replicates from two independent experiments).

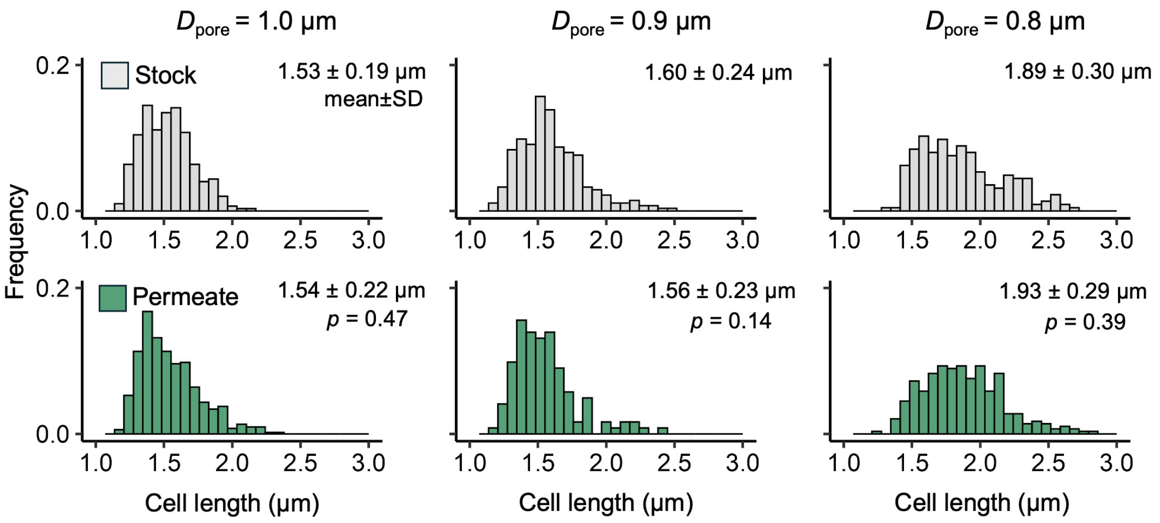


**Figure S12.** Cell-length distributions for *E. coli* in the starting population and among cells recovered in the permeate following mechanofiltration through filters with pore diameters of 1.0, 0.9, and 0.8 µm. Cell length does not differ between stocks and permeates, suggesting that length variation does not influence cell transport. Data include >100 cells per condition collected from 3–4 permeates and compared with time-matched stock populations. Statistical comparisons in were performed using Dunnett's test versus the stock population.

**Table S1:** Conversion of cell stiffness ($k$) differences reported in literature used to calculate fold-changes in cell compliance.

| Condition | Fold-change stiffness $\left(\frac{k_{condition}}{k_{untreated,\ wild-type}}\right)$ | Fold-change compliance $\left(\frac{k_{condition}}{k_{untreated,\ wild-type}}\right)^{-1}$ | Reference |
|---|---|---|---|
| Δ*ompA* | 0.57 | 1.8 | Rojas, *Nature* 2018 |
| EDTA | 0.62 | 1.6 | Rojas, *Nature* 2018 |
| A22 | 0.50 | 2.0 | Wang, *PNAS* 2010 |
| Δ*lpp* | 0.41 | 2.4 | Rojas, *Nature* 2018 |

**SUPPLEMENTAL TEXT – THEORY**

**Estimation of Hydrostatic Pressure:**

The magnitude of hydrostatic pressure (*P*) generated during centrifugation in mechanofiltration is described by **Eqn. 1**:

$$P = \omega^2 \rho \left( r_o h + \frac{h^2}{2} \right), \qquad \text{(Eqn. 1)}$$

where, ω represents the angular velocity, $\rho$ is equal to the density of liquid within the insert (assumed equal to water at 1 g/mL), $r_o$ is the distance between the center of the centrifuge and top of liquid within the well, and $h$ is the height of the liquid column above the filter. We assume hydrostatic pressure is distributed equally across the surface of the filter. Thus, the force from hydrostatic pressure that drives cell transport through pores ($F_{\text{hydrostatic}}$) is the same across all mechanofiltration experiments if apical well volume and centrifugation parameters are conserved.

*Significance and Limitations:* **Eqn. 1** assumes a constant angular velocity. However, during centrifugation, the angular velocity ramps to a maximum, is maintained constant for a period of time, and then decreases. Additionally, the angular velocity at which liquid efflux from the apical chamber occurs is not known. In our experiments the maximum angular velocity is 4000 rpm, and the centrifuge radius $r_0$ is 13.4 cm. Unless specified, we added 200 µL of bacterial suspension to the apical chamber of the well at the start of the experiment, meaning that $h$ was equal to 6 mm, based on insert dimensions. Thus, a maximum of 139 kPa of hydrostatic could be generated. To alter loading conditions, we reduced apical well volume to 100 µL, reducing $h$ to 3 mm and maximum pressure to 70 kPa.

We assume that during centrifugation, multiwell plates are positioned completely perpendicular to the rotor arm. However, there likely exists a small angle between the plate and centrifuge when swinging bucket rotors are used. This angle will influence hydrostatic pressure by changing $r_0$ across the plate. For example, a theoretical pitch angle of 10° would lead to a 10% reduction in pressure between wells at the top and bottom of the plate. To mitigate the potential effects of misalignment, we performed all experiments reported in the main text within wells positioned at the center of the plate (rows: B-E, columns: 3-6).

**Derivation of Analytical Model to Describe Cell Transport through Pores**

At a given pore size, the force from hydrostatic pressure will deform bacteria into and through pores until the difference between cell diameter ($D_{\text{cell}}$) and pore diameter ($D_{\text{pore}}$) becomes too large. We define this limit as the maximum deformation ($\Delta_{\max}$). To understand how $\Delta_{\max}$ relates to the material properties of the cell, we consider that the entry of a bacterium into a pore that is narrower than cell width is similar to the loading of an oversized shaft into an undersized hub, which occurs in 'press fit' components. Forces associated with press fit assembly are dependent on the interfacial pressure ($P_{\text{i}}$) between the shaft and hub when assembled[1]. $P_{\text{i}}$ can be determined using Lame's Equation as follows:

$$P_i = \frac{\delta}{D} \cdot \frac{1}{\frac{1}{E_{hub}}\left(\frac{d_o^2+D^2}{d_o^2-D^2}+\upsilon_{hub}\right)+\frac{1}{E_{shaft}}\left(\frac{D^2+d_i^2}{D^2+d_i^2}-\upsilon_{shaft}\right)}\ , \qquad \text{(Eqn. 2)}$$

where δ is the difference in diameters between the shaft and the hole, $E$ describes the Young's modulus of the shaft or hub material, υ Poisson's ratios, and $D$ the shared diameter of the hub and shaft. Conventionally, the shaft and hub are modeled as hollow, thick-walled cylinders with inner diameter of $d_i$ for the shaft and outer diameter of $d_o$ for the hub. To adapt the press fit theorem to mechanofiltration, we instead represent the cell as a solid shaft so that $d_i$ equals 0, and we model the filter pore as a hub with infinite wall thickness ($d_o \rightarrow \infty$). **Eqn. 2** thus becomes:

$$P_i = \frac{\delta}{D_{pore}} \cdot \frac{1}{\frac{1}{E_{filter}}(1+ \upsilon_{filter})+\frac{1}{E_{cell}}(1- \upsilon_{cell})} \quad , \qquad \text{(Eqn. 3)}$$

where, the shared diameter ($D$), becomes $D_{\text{pore}}$ if we assume that the pore does not expand significantly upon cell entry. To facilitate cell entry and transport through the pore in mechanofiltration, the downward force from hydrostatic pressure ($F_{\text{hydrostatic}}$), must be greater than the force from $P_{\text{i}}$ times the coefficient of friction, $\mu$, between the cell and pore (Coulomb's Law). $P_{\text{i}}$ acts over the contact area between the cell and pore walls, which is equal to the surface area of the cell trunk within the pore ($\pi D_{\text{pore}} C_{\text{Length}}$). Thus, the greatest deformation ($\Delta_{\text{max}}$) that can be achieved by cells under $F_{\text{hydrostatic}}$ is:

$$\Delta_{max} = \frac{F_{hydrostatic}}{\pi C_{Length} \mu} \left( \frac{1}{E_{filter}} \left(1 + \upsilon_{filter}\right) + \frac{1}{E_{cell}} \left(1 - \upsilon_{cell}\right) \right), \qquad \text{(Eqn. 5)}$$

*Model Implications and Agreement with Experimental Results:* In agreement with our experimental results, **Eqn. 5** indicates that $\Delta_{\text{max}}$ increases linearly with $F_{\text{hydrostatic}}$ and decreases as cell stiffness ($E_{\text{cell}}$) increases. $\Delta_{\text{max}}$ also depends on $C_{\text{Length}}$; however, minor variation in $C_{\text{Length}}$ observed within bacterial populations did not affect the probability of cell transport (**Figure S12**). Future work is needed to determine whether larger variation in $C_{\text{Length}}$ will influence transport. $\Delta_{\text{max}}$ will also be influenced by a cell's Poisson's ratio. It is possible that the genetic and chemical perturbations made to the cell envelope to reduce whole-cell stiffness also affect the Poisson's ratio of the cell and contribute to our observed changes in $\Delta_{max}$; these effects remain to be investigated. Finally, **Eqn. 5** suggests that $\Delta_{max}$ will also be dependent upon the material properties of the filter. All filters used in this study were made from the same track-etched, polyester membrane, and thus variations in $E_{filter}$ or $\upsilon_{filter}$ did not affect our results.
